# *Plasmodium falciparum* has evolved multiple mechanisms to hijack human immunoglobulin M

**DOI:** 10.1101/2022.08.03.502706

**Authors:** Chenggong Ji, Hao Shen, Chen Su, Yaxin Li, Shihua Chen, Thomas H. Sharp, Junyu Xiao

## Abstract

*Plasmodium falciparum* causes the most severe malaria in humans. Immunoglobulin M (IgM) serves as the first line of humoral defense against infection and potently activates the complement pathway to facilitate *P. falciparum* clearance. A number of *P. falciparum* proteins hijack IgM, leading to immune evasion and severe disease. However, the underlying molecular mechanisms remain unknown. Here, using high-resolution cryo-electron microscopy, we delineate how *P. falciparum* proteins VAR2CSA, TM284VAR1, DBLMSP, and DBLMSP2 target IgM. Each protein binds IgM in a different manner, and together they present a variety of Duffy-binding-like domain-IgM interaction modes. We further show that these proteins interfere directly with IgM-mediated complement activation, with VAR2CSA exhibiting the most potent inhibitory effect. Structural analyses suggest that VAR2CSA occludes the congregation of the complement C1 complex on IgM. These results underscore the importance of IgM for the adaptation of *P. falciparum* to humans, and provide critical insights into the immune evasion mechanism of *P. falciparum*.

## Introduction

Malaria is one of the greatest killers of humankind throughout history and remains a major public health problem: approximately 241 million malaria cases were documented worldwide in 2020, resulting in 627,000 deaths ^1^. Malaria is caused by infection with *Plasmodium* parasites, among which *Plasmodium falciparum* causes the most devastating disease. The merozoite form of *P. falciparum* invades the red blood cells to replicate inside, and the infected red blood cells (iRBCs) are eventually ruptured to release more merozoites, resulting in fever and hemolytic anemia. Furthermore, iRBCs can adhere to the placenta and brain endothelium, leading to fatal complications known as placental and cerebral malaria.

Immunoglobulins, or antibodies, are central components of the immune system and provide critical protections against various pathogens, including *P. falciparum*. The immunoglobulin M (IgM) type of antibodies is the first to be produced in a humoral immune response ^2,3^. The predominant form of IgM is an asymmetrical pentamer, with five IgM monomers joined together by the joining chain (J-chain) ^4,5^. The presence of ten antigen-binding sites within an IgM pentamer allows it to bind and neutralize pathogens effectively. Furthermore, IgM efficiently activates the complement pathway, which plays a crucial role in malaria immunity ^6^.

During the evolutionary arms race between the *Plasmodium* parasite and humankind, *P. falciparum* has evolved strategies to antagonize the function of IgM. *Plasmodium falciparum* erythrocyte membrane protein 1 (PfEMP1) is a family of ~60 virulent proteins secreted by *P. falciparum* to the iRBC surface. PfEMP1 proteins have very large extracellular segments, consisting of different numbers and types of Duffy-binding-like (DBL) domains and cysteine-rich interdomain regions. These versatile modules endow PfEMP1 proteins with the ability to interact with a range of molecules in humans ^7–9^. For example, VAR2CSA, a member of the PfEMP1 family and a major culprit in placental malaria, can bind to chondroitin sulfate A (CSA) glycosaminoglycans, resulting in the sequestration of iRBCs within the placenta ^10^. For this reason, VAR2CSA is considered a prominent target for the development of placental malaria vaccines. VAR2CSA also interacts with IgM, and appears to employ IgM as a shield to conceal itself from immunoglobulin G (IgG) antibodies ^11,12^. Similarly, a number of other PfEMP1 variants bind to IgM ^13–15^, and the presence of nonimmune IgM on iRBCs correlates with severe malaria ^16^. In addition, two other *P. falciparum* proteins, DBLMSP and DBLMSP2, are capable of hijacking IgM ^17^. In contrast to the PfEMP1 proteins that reside on iRBCs, DBLMSP and DBLMSP2 are located on the surface of *P. falciparum* merozoites ^18^, which are directly exposed to humoral immunity. It is likely that the nonimmune IgM also provides camouflage for merozoites and facilitates their evasion of IgG antibodies ^17^.

## Results

### Cryo-EM structure determination

To understand how these *P. falciparum* proteins specifically hijack IgM, we prepared the ectodomains of two PfEMP1 proteins, VAR2CSA (from the FCR3 strain) and TM284VAR1 (from a cerebral parasite strain ^19^); as well as the DBL domains of DBLMSP (from field isolate 017 ^17^) and DBLMSP2 (from the 3D7 strain) (Fig. 1a), and tested their interactions with the human pentameric IgM core (Fcμ-J) that consists of the-IgM-Fc (Fcμ) pentamer and the J-chain ^4^. Surface plasmon resonance (SPR) analyses demonstrate that each *P. falciparum* protein binds to Fcμ-J with high affinity, exhibiting *K*_d_ values of 7-30 nM (Fig. 1b).

**Fig. 1.**
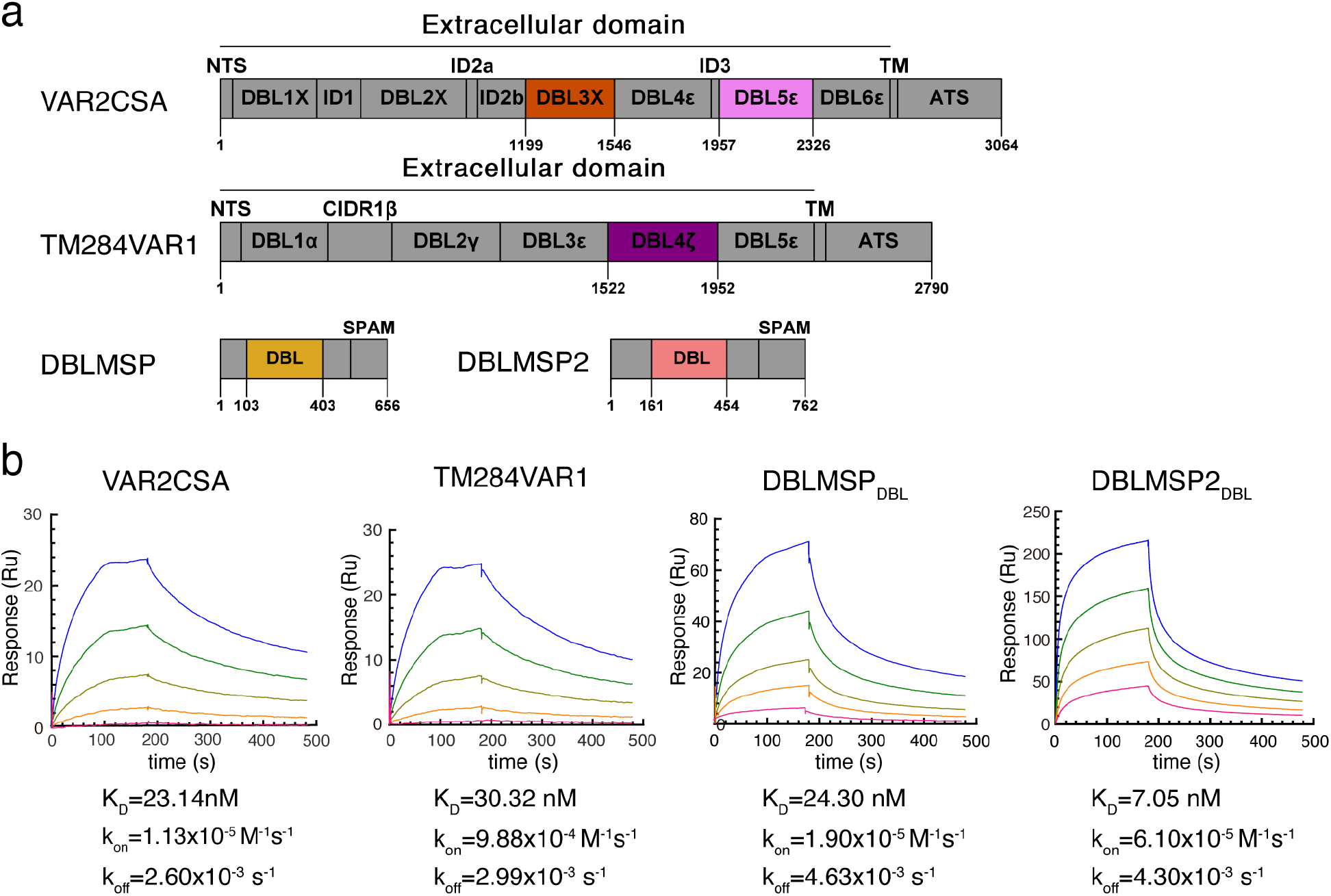
The *P. falciparum* proteins directly interact with the human pentameric IgM core. **a.** Schematics of the domain organizations of VAR2CSA, TM284VAR1, DBLMSP, and DBLMSP2. **b.** Surface plasma resonance analyses of the interactions between the *P. falciparum* proteins and Fcμ-J.

We subsequently reconstituted their complexes with Fcμ-J (Extended Data Fig. 1) and determined the cryo-electron microscopy (cryo-EM) structures of these complexes (Fig. 2, Extended Data Fig. 2–3, Extended Data Table 1). Although some PfEMP1 proteins can bind IgM in a 2:1 ratio ^20,21^, only 1:1 complexes were observed in our study. Fcμ-J exhibits an pentameric architecture, with the J-chain conferring asymmetry on the central Fcμ platform, as seen in the complex with the secretory component (SC), i.e., the ectodomain of the polymeric immunoglobulin receptor (pIgR) ^4,5^. The pIgR/SC-binding face of Fcμ-J is also targeted by the *P. falciparum* proteins (Extended Data Fig. 4), and the interactions between the *P. falciparum* proteins and Fcμ-J exclusively involve the Fcμ-Cμ4 domains, which is consistent with previous analyses ^11,17,19^. The DBL domains in the *P. falciparum* proteins are responsible for interacting with Fcμ; strikingly however, they display distinct Fcμ-binding modes.

**Fig. 2.**
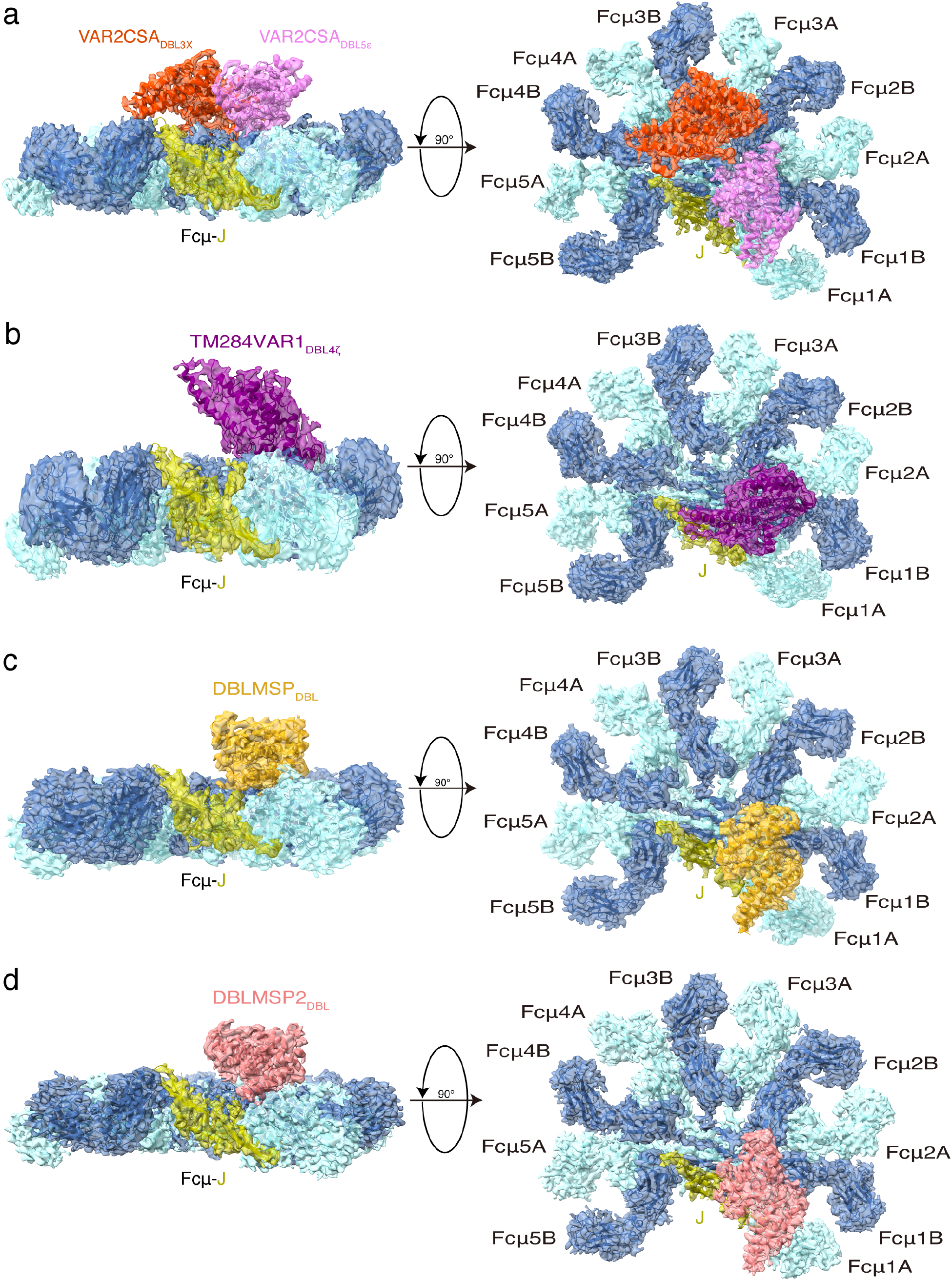
Cryo-EM structures of the *P. falciparum* proteins in complex with the human pentameric IgM core. **a.** Structure of the VAR2CSA-Fcμ-J complex. Only the DBL3X and DBL5ε domains of VAR2CSA are shown. **b.** Structure of the TM284VAR1_DBL4ζ_-Fcμ-J complex. **c.** Structure of the DBLMSP_DBL_-Fcμ-J complex. **d.** Structure of the DBLMSP2_DBL_-Fcμ-J complex.

### Structure of the VAR2CSA-Fcμ-J complex

The ~300 kDa ectodomain of VAR2CSA consists of six DBL domains plus the interdomain regions (IDs) (Fig. 1a). Recent cryo-EM studies demonstrated that the regions encompassing DBL2X-ID3 assemble into a stable core, whereas DBL5ε-DBL6ε forms a flexible arm ^22–24^. The majority of this large molecule can be visualized in the density map of the VAR2CSA-Fcμ-J complex (Extended Data Fig. 2c). The IgM-binding sites in VAR2CSA have previously been variously mapped to DBL2X, DBL5ε, and DBL6ε ^25,26^; however, our cryo-EM structure unambiguously reveals that DBL3X and DBL5ε conjointly mediate the binding to Fcμ (Fig. 2a). This result is highly concordant with previous observations showing that IgM specifically excludes the binding of DBL3X- or DBL5ε-specific IgGs to RBCs infected by VAR2CSA-expressing *P. falciparum* parasites ^11,27^. The major CSA binding pocket is formed by several domains within the stable core of VAR2CSA, especially DBL2X and DBL4ε. DBL3X and DBL5ε are located distal to the CSA-binding site; therefore VAR2CSA should be able to bind to IgM and CSA simultaneously (Fig. 3a). Indeed, previous studies showed that IgM did not affect the adhesion of VAR2CSA-bearing iRBCs to CSA ^11^.

**Fig. 3.**
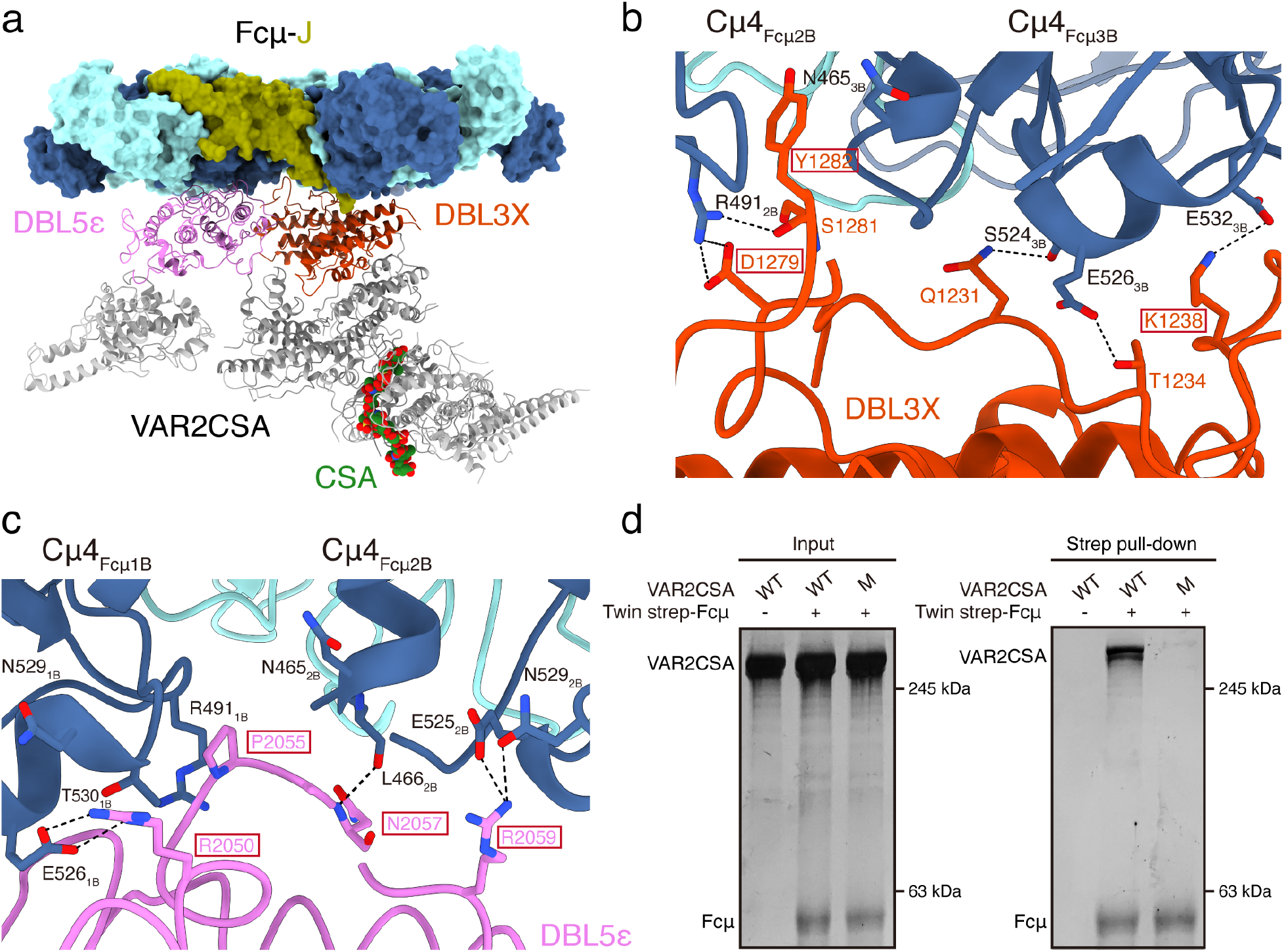
VAR2CSA targets Fcμ via DBL3X and DBL5ε and inhibits IgM-mediated complement activation. **a.** A composite structural model of VAR2CSA binding to both Fcμ-J and CSA. **b.** VAR2CSA_DBL3X_ interacts with Cμ4 in Fcμ2B and Fcμ3B via subdomain SD1. VAR2CSA residues that are mutated in VAR2CSA-M are highlighted by red boxes. **c.** VAR2CSA_DBL5ε_ interacts with Fcμ1B and Fcμ2B. **d.** VAR2CSA-M displays reduced binding to Fcμ-J.

DBL3X and DBL5ε together interact with three Fcμ units within the Fcμ pentamer. Both DBL3X and DBL5ε exhibit an archetypical DBL fold that can be further divided into three subdomains ^28,29^: SD1 comprises mostly loops, whereas SD2 and SD3 contain a characteristic four-helix bundle and double-helix hairpin, respectively (Extended Data Fig. 5). DBL3X interacts with the Cμ4 domains of Fcμ2B (the ten Fcμ chains in the Fcμ pentamer are named as previously described, starting from the Fcμ chain that interacts with the C-terminal hairpin of the J-chain as 1A ^4^) and Fcμ3B using residues in subdomain SD1 (Fig. 3b). Tyr1282 is sandwiched between Fcμ2B and Fcμ3B and packs against Arg491_Fcμ2B_ and Asn465_Fcμ3B_. Arg491_Fcμ2B_ is also contacted by Asp1279, whereas Lys1238 forms an ion pair with Glu532_Fcμ3B_. DBL5ε, on the other hand, interacts with Fcμ1B and Fcμ2B, also mainly using a loop in SD1 (Fig. 3c). Pro2055-Ala2056 insert between Fcμ1B and Fcμ2B and pack with Arg491_Fcμ1B_ and Asn465_Fcμ2B_. Arg2050 packs with Asn529_Fcμ1B_-Thr530_Fcμ1B_, and also coordinates Glu526_Fcμ1B_. Arg2059 interacts with Glu525_Fcμ2B_ and Asn529_Fcμ2B_. A VAR2CSA heptamutant (VAR2CSA-M), K1238A/D1279A/Y1282A/R2050A/P2055G/N2057A/R2059A, failed to interact with Fcμ-J, validating the functional relevance of the molecular interactions described above (Fig. 3d).

### VAR2CSA directly inhibits IgM-mediated complement-dependent cytotoxicity

IgM is a potent activator of the classical complement pathway; however, it has long been documented that the recruitment of IgM onto iRBCs by VAR2CSA does not render iRBCs susceptible to complement-dependent cytotoxicity (CDC) ^11^. In fact, another PfEMP1 protein, IT4VAR60, binds to IgM and blocks the deposition of C1q (a key component of the complement C1 complex) on the iRBCs, thereby protecting the iRBCs from complement-mediated lysis ^21^. To examine whether VAR2CSA directly inhibits CDC, we prepared a recombinant anti-CD20 IgM molecule by engineering the antigen-binding fragment of rituximab, a monoclonal antibody against CD20, onto Fcμ (Extended Data Fig. 1f). In the presence of human serum complement, this IgM molecule robustly triggered the lysis of CD20+ OCI-Ly10 cells (Extended Data Fig. 1g). Preincubation of this IgM with VAR2CSA greatly reduced its ability to activate CDC, with a half maximal inhibitory concentration (IC_50_) of ~1.9 nM (Fig. 4a). In contrast, VAR2CSA-M displayed no such inhibitory effect, further corroborating the above structural and biochemical analyses.

**Fig. 4.**
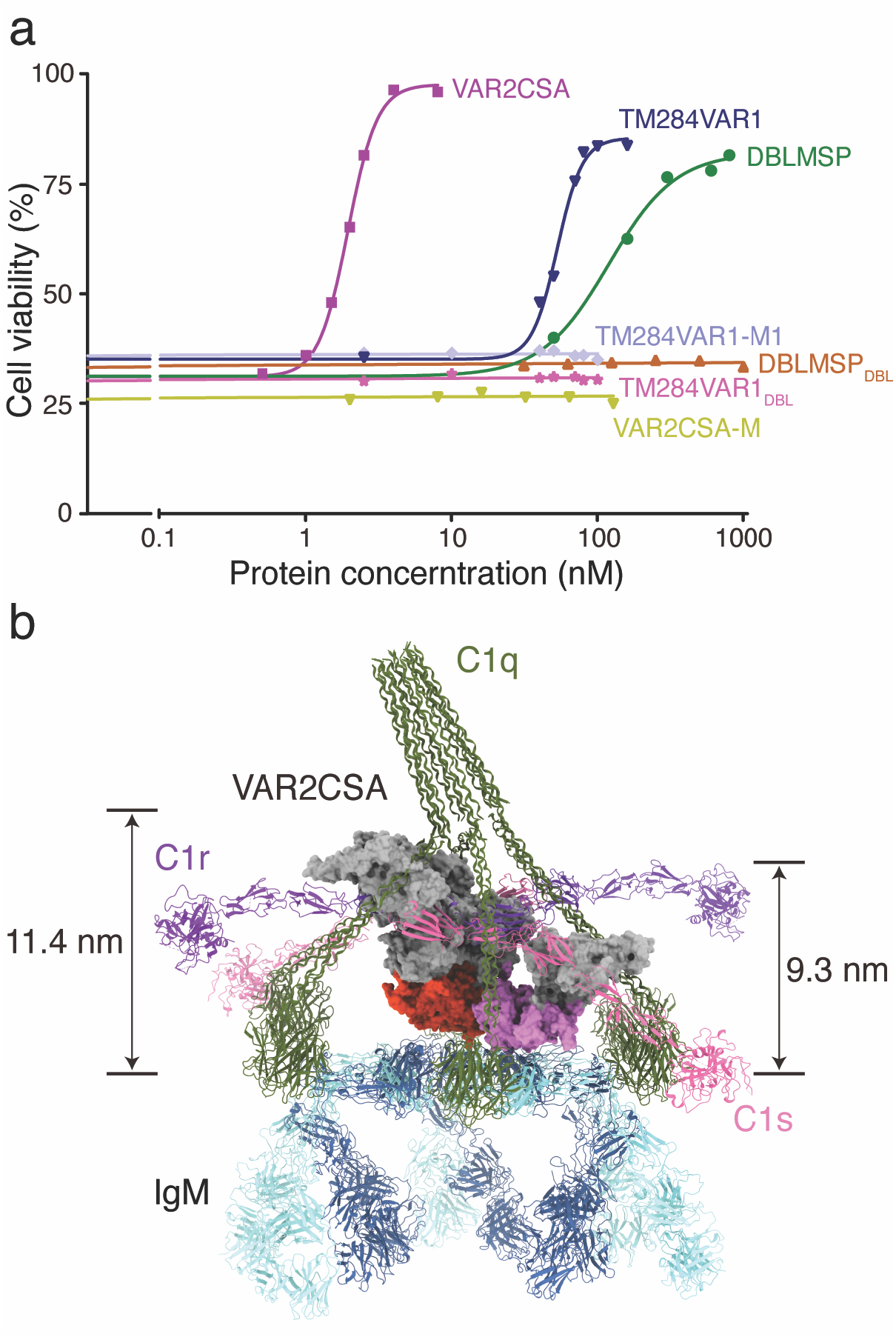
The *P. falciparum* proteins directly inhibit complement dependent cytotoxicity. **a.** VAR2CSA, TM284VAR1, and DBLMSP all inhibit IgM-mediated CDC. **b.** VAR2CSA inhibits CDC by sterically occluding the interaction between IgM and C1.

It has been suggested that IT4VAR60 occupies the C1q binding site on IgM ^21^. The C1q binding site is located in the Cμ3 domain of Fcμ and critically involves residues 432-436 in the FG loop ^30^, which are positioned at the outer edge of Cμ3. The binding region of VAR2CSA is confined to the central Cμ4 domains and does not extend to the C1q binding site. However, it appears that VAR2CSA can block the binding of C1 to IgM via steric hindrance (Fig. 4b). The central cavity of the C1 complex that accommodates the C1r/C1s proteases has a height of ~9.3 nm above the Fcμ platform ^31^, which is surpassed by the ~11.4 nm stature of VAR2CSA. Thus, VAR2CSA, with its colossal size, would effectively occlude the congregation of the C1 complex on IgM and thus directly blocks the initiation of CDC.

### TM284VAR1_DBL4ζ_ is responsible for interacting with Fcμ-J

The ectodomain of TM284VAR1 exhibits a flexible structure, and a 3D reconstruction at ~8 Å reveals an elongated architecture that somewhat resembles that of VAR2CSA, with a length of ~14 nm (Extended Data Fig. 6). Although the entire ectodomain was used to reconstitute the complex with Fcμ-J (Extended Data Fig. 1b), only the DBL4ζ domain can be clearly visualized in the density map (Fig. 2b, Extended Data Fig. 2i). Other regions likely display conformational disorder and are not discernible after single-particle averaging. This is also highly concordant with previous biochemical analyses demonstrating that the DBL4ζ domain in TM284VAR1 is solely responsible for binding to IgM ^19,32^.

DBL4ζ interacts with Fcμ1B, Fcμ2A, and Fcμ2B using residues from both SD1 and SD2. In particular, the α3 helix and the following loop within the SD2 four-helix bundle provide a focal point for the DBL4ζ-Fcμ interaction (Fig. 5a). Because of this tight interaction, this region displays high-quality density with clear side chain features (Extended Data Fig. 2n). Glu1705 binds with Arg491_Fcμ1B_. Arg1706 forms a bidentate interaction with Glu468_Fcμ2B_. Lys1709 and Arg1712 interact with Asp453_Fcμ2A_. Asp1716 and Asn1717 interact with the main chain groups of Ala542_Fcμ2A_ and Leu449_Fcμ2A_. Two TM284VAR1 mutants, E1705A/R1706A/K1709A (M1) and E1705A/R1706A/D1716A (M2), display diminished interactions with Fcμ-J (Fig. 5b), confirming the critical functions of these residues in binding to IgM. Furthermore, the Tyr1728-Tyr1732 loop packs intimately with the Gly492_Fcμ1B_-Pro497_Fcμ1B_ loop (Fig. 5c). Indeed, bovine and mouse IgM differ significantly from human IgM at the Gly492-Pro497 loop, and neither of them binds to TM284VAR1 ^19,26^. Additionally, the replacement of the human IgM residues Pro494-Pro497 with the corresponding mouse sequence also abolished the interactions with TM284VAR1_DBL4ζ_ ^19^. Similar to VAR2CSA, the ectodomain of TM284VAR1 suppressed IgM-mediated complement activation, although not as potently, displaying an IC_50_ of ~52.9 nM (Fig. 4a). In contrast, neither TM284VAR1-M1 nor TM284VAR1_DBL4ζ_ exerted such an effect. It is likely that TM284VAR1 also interferes sterically with the IgM-C1 interaction, and the relatively mild inhibition is caused by its less rigid structure compared to that of VAR2CSA.

**Fig. 5.**
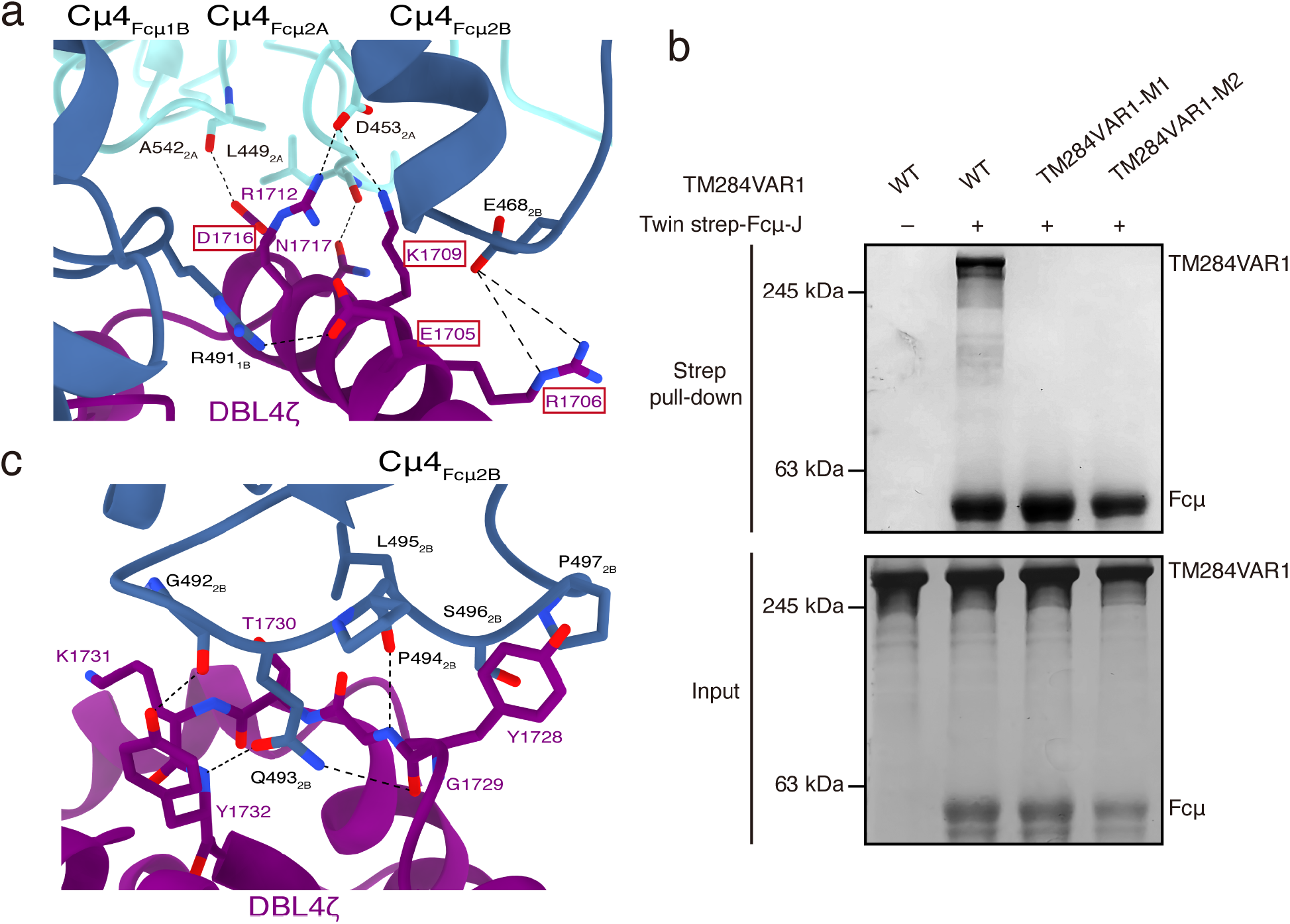
TM284VAR1 interacts with Fcμ using the DBL4ζ domain. **a.** Interactions between TM284VAR1_DBL4ζ_ and Fcμ. **b.** TM284VAR1 mutants display reduced binding to Fcμ-J. **c.** Interactions between the TM284VAR1_DBL4ζ_ Tyr1728-Tyr1732 loop and Fcμ.

### DBLMSP_DBL_ and DBLMSP2_DBL_ display another Fcμ-binding mode

DBLMSP comprises one DBL domain that is responsible for interacting with IgM ^17^ and a SPAM domain that is involved in oligomerization ^33^. As the full-length DBLMSP protein is unstable and tends to form heterogeneous oligomers in solution (Extended Data Fig. 1c), we performed a cryo-EM study using only the DBL domain. The resulting structure shows that DBLMSP_DBL_ interacts intimately with Fcμ1A, Fcμ1B, and Fcμ2B (Fig. 2c) using all three subdomains. In SD1, a loop involving DBLMSP residues Ile140-Ala143 lodges in the groove between Cμ4_Fcμ1A_ and Cμ4_Fcμ1B_, whereas His173-Arg174 contact Cμ4_Fcμ2B_ (Fig. 6a). In SD2, Asp229-Ile232 pack with the 525-530 helix in Cμ4_Fcμ1B_ (Fig. 6b). Glu352, Asn356, and Arg357 in SD3 engage Cμ4_Fcμ1A_ (Fig. 6c). Indeed, two DBLMSP_DBL_ mutants, N169A/H173A/R174A and D229A/Y230A/Q231A, do not bind to Fcμ-J (Fig. 6d). Interestingly, oligomeric full-length DBLMSP interferes with complement activation by IgM (IC_50_ ~119.1 nM, which could be an underestimate due to the unstable nature of recombinant full-length DBLMSP); whereas the monomeric DBLMSP_DBL_ domain is not capable of doing so (Fig. 4a). This result further supports our hypothesis that the large PfEMP1 proteins or DBLMSP oligomer antagonize IgM-mediated complement activation via steric hindrance.

**Fig. 6.**
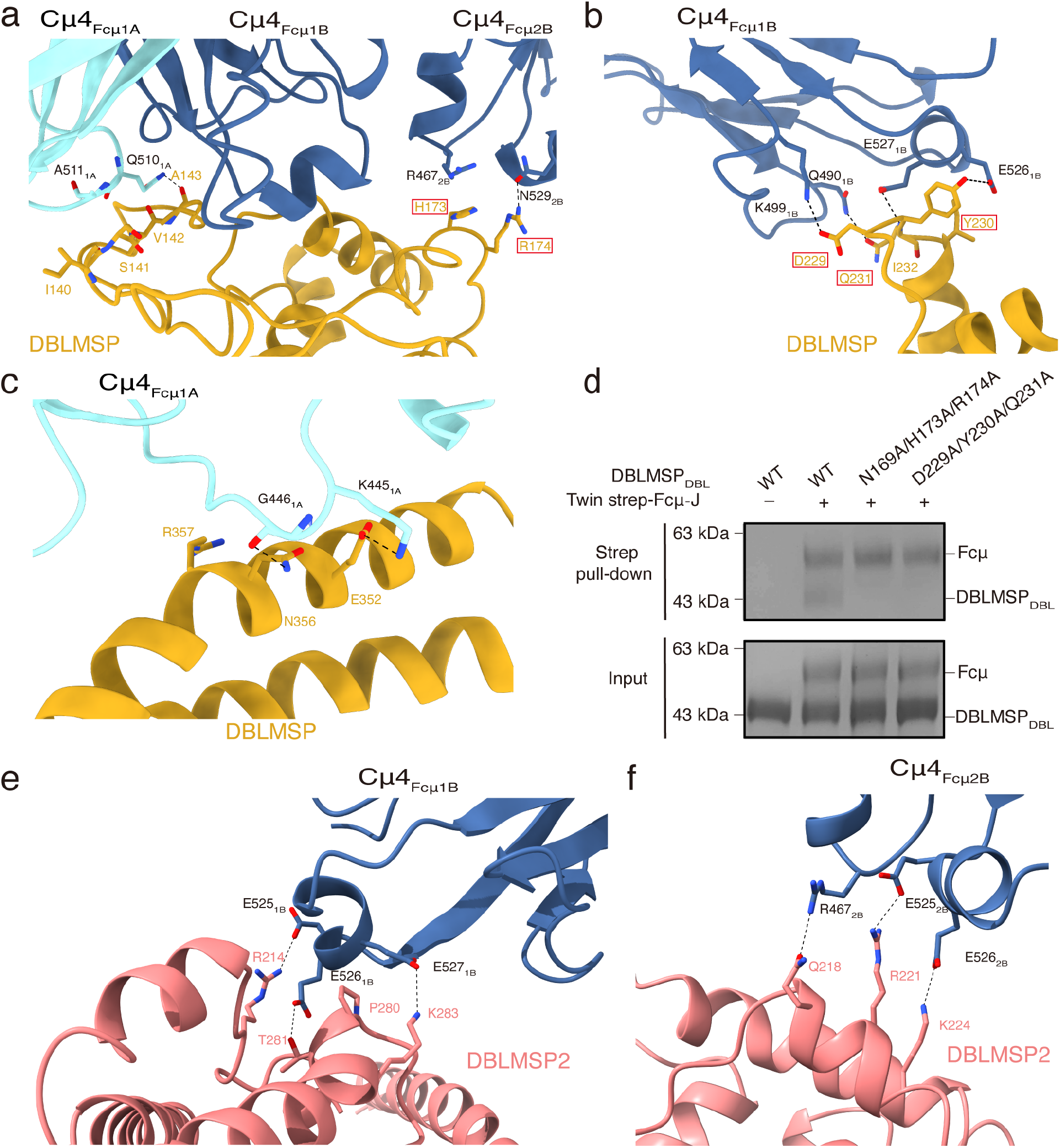
Interactions between DBLMSP family proteins and Fcμ. **a.** Interactions between DBLMSP-SD1 and Fcμ. **b.** Interactions between DBLMSP-SD2 and Fcμ. **c.** Interactions between DBLMSP-SD3 and Fcμ. **d.** DBLMSP_DBL_ mutants display reduced binding to Fcμ-J. **e.** DBLMSP2_DBL_ form extensive interactions with three consecutive Glu in Cμ4_Fcμ1B_. **f.** DBLMSP2_DBL_ interacts with Glu525-Glu526 in Cμ4_Fcμ2B_.

We also determined the cryo-EM structure of the DBLMSP2_DBL_-Fcμ-J complex. When compared to the DBLMSP_DBL_-Fcμ-J structure, an overall similar pattern of interaction between DBLMSP2_DBL_ and Fcμ-J is observed (Fig. 2c, 2d). Similar to DBLMSP_DBL_, DBLMSP2_DBL_ also targets Fcμ1A, Fcμ1B, and Fcμ2B; and all three of its subdomains are involved in binding to these Fcμ molecules. Distinct molecular interactions are nevertheless present at the binding interfaces. For example, several DBLMSP2 residues, including Arg214 from SD1, as well as Pro280, Thr281, and Lys283 from SD2, form extensive ionic, van der Waals, and hydrogen bond interactions with three consecutive Glu in Cμ4_Fcμ1B_ (Glu525-Glu527, Fig. 6e). Arg221 and Lys224 contact Glu525-Glu526 in Cμ4_Fcμ2B_ (Fig. 6f). These interactions are unique to the DBLMSP2_DBL_-Fcμ-J complex, and likely contribute to the high binding affinity between DBLMSP2_DBL_ and Fcμ-J (Fig. 1b).

## Discussion

IgM serves as the first line of defense in adaptive immunity and initiates the complement cascade to communicate with the innate immune system. The ability to hijack IgM bestows a survival advantage on *P. falciparum* and therefore can increase virulence. The four *P. falciparum* proteins we investigated all target the Cμ4 domains of Fcμ using their DBL domains; together, however, they show a variety of Fcμ-binding modes (Fig. 7). VAR2CSA_DBL5ε_, TM284VAR1_DBL4ζ_, DBLMSP_DBL_, and DBLMSP2_DBL_ all interact with Fcμ1-Fcμ2, and their binding sites overlap with those of pIgR/SC (Extended Data Fig. 4). Nevertheless, the molecular interactions between these three DBL domains and Fcμ1-Fcμ2 are different. VAR2CSA_DBL5ε_ covers the base of Fcμ1B and Fcμ2B using subdomain SD1, and the remaining SD2-SD3 subdomains are projected toward Fcμ1. TM284VAR1_DBL4ζ_ mainly binds to Fcμ1-Fcμ2 using subdomain SD2 and approaches the Fcμ plane using a completely different angle. All three subdomains of DBLMSP_DBL_ and DBLMSP2_DBL_ are involved in interacting with Fcμ1-Fcμ2. In contrast to these DBL domains that target Fcμ1-Fcμ2, VAR2CSA_DBL3X_ engages Fcμ2-Fcμ3 instead, and also uses a different set of SD1 residues from VAR2CSA_DBL5ε_ to target these Fcμ molecules (Extended Data Fig. 5). The DBL domains display high sequence diversity and are adaptable, interacting with a myriad of molecules ^34^. It is amazing how they have evolved such diverse ways to target one human molecule. This is reminiscent of the two binding modes between the DBL domains and ICAM-1 ^35^ and truly underscores the paramount importance of IgM for malaria immunity. It is noteworthy that all these DBL domains interact with multiple Fcμ molecules within the Fcμ pentamer; therefore, it is unlikely that they will target the monomeric form of IgM, as in the B-cell receptor complex. In fact, VAR2CSA_DBL3X_, VAR2CSA_DBL5ε_, and TM284VAR1_DBL4ζ_ all heavily exploit the space between adjacent Fcμ molecules for binding. On the other hand, none of them form significant interactions with the J-chain, so it is likely that they will bind equally well to an IgM hexamer, which is devoid of the J-chain.

**Fig. 7.**
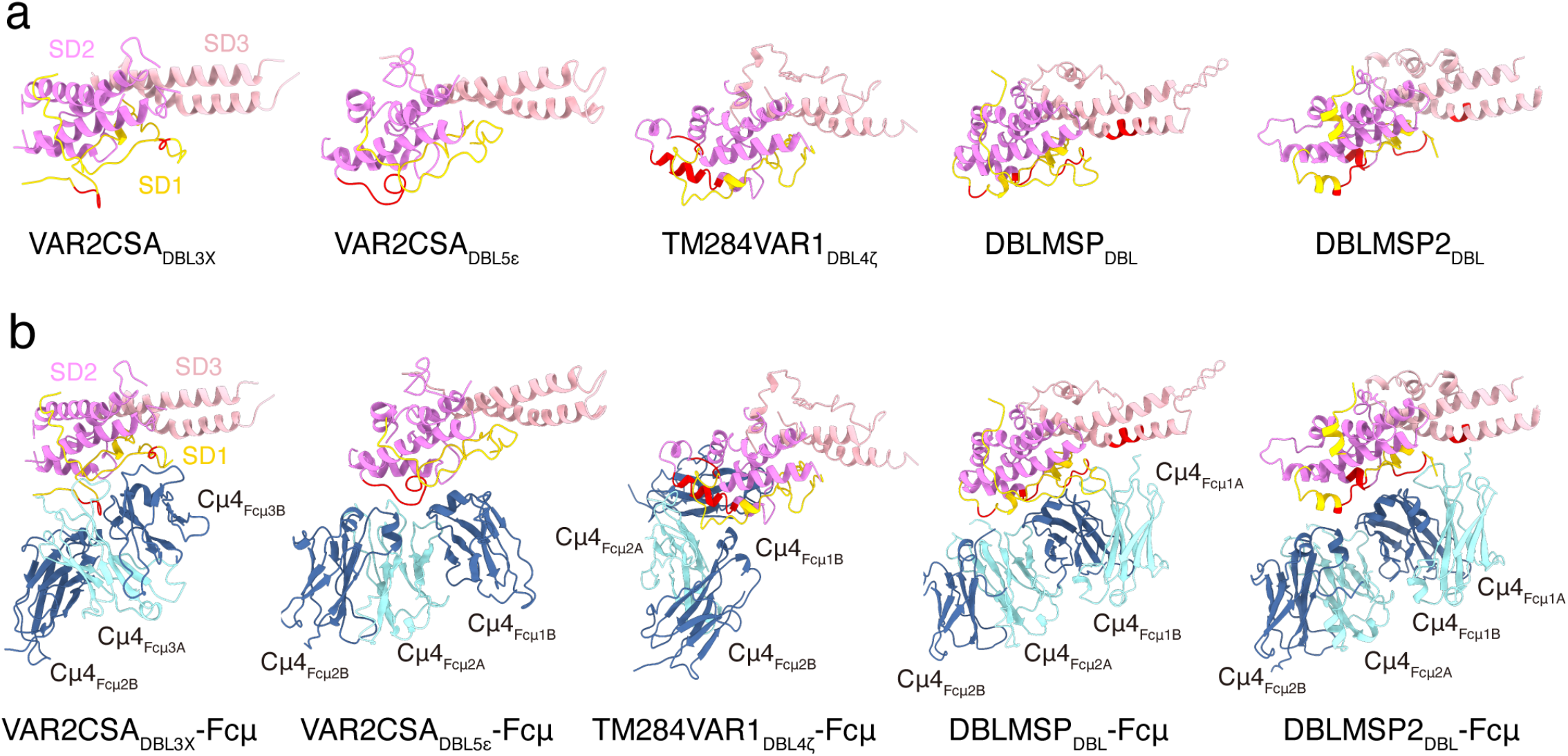
Different Fcμ-binding modes. **a.** VAR2CSA_DBL3X_, VAR2CSA_DBL5ε_, TM284VAR1_DBL4ζ_, DBLMSP_DBL_, and DBLMSP2_DBL_ are shown in the same orientation, with their SD1, SD2, and SD3 subdomains colored yellow, purple, and pink, respectively. The regions involved in binding to Fcμ are highlighted in red. **b.** VAR2CSA_DBL3X_, VAR2CSA_DBL5ε_, TM284VAR1_DBL4ζ_, DBLMSP_DBL_, and DBLMSP2_DBL_ in complexes with the Fcμ molecules.

How does the hijacking of IgM by these proteins benefit *P. falciparum* parasites? The main goal appears to be immune evasion ^36^. First, immune IgM plays a critical role in malaria immunity ^6,37^, and these proteins can demobilize immune IgM molecules by grabbing Fcμ, thereby preventing them from binding to their cognate targets during *P. falciparum* infection. Second, IgM facilitates the masking of antibody epitopes. A number of antibody epitopes have been identified and mapped onto the structure of VAR2CSA ^22,27,38–40^. Apparently, most of the epitopes in DBL3X and DBL5ε can be concealed by Fcμ-J, such as PAM8.1, P62, P63, and P23 (Extended Data Fig. 7). Notably, Fcμ-J only represents the IgM core. An entire IgM molecule, with Cμ2 and also the antigen-binding fragments, can have a length of ~38 nm in its fully extended conformation. This large size makes an ideal umbrella for the *P. falciparum* parasites to block the attack of neutralizing antibodies. Third, these proteins can directly interfere with IgM-mediated complement activation (Fig. 4a), thereby disarming another important force in human immunity. Finally, these proteins can impede the interaction between IgM and its cellular receptors. As described above, all these proteins dwell in the interaction “hot spot” of IgM and clearly interfere with the binding of pIgR/SC (Extended Data Fig. 4), which governs the mucosal transport of IgM. The other two IgM receptors, FcαμR and FcμR, function in the humoral immune response and are likely more relevant for malaria immunity. The molecular mechanisms of how these two receptors engage IgM are less understood; however, the Cμ4 domain of IgM is critical for their binding ^41–43^. Therefore, the *P. falciparum* proteins could interfere with the perception of IgM by its receptors and suppress IgM-related immune signaling pathways.

## Methods

### Cell culture

Sf21 and High Five insect cells were cultured using SIM-SF and SIM-HF media (Sino Biological) in a nonhumidified shaker at 27 °C. HEK293F cells were cultured using SMM 293-TI medium (Sino Biological) in a humidified shaker at 37 °C with 5% CO_2_. OCI-Ly10 cells were cultured using RPMI-1640 (Gibco) medium supplemented with 10% fetal bovine serum (Gibco) and 1% penicillin-streptomycin (Gibco) in a humidified incubator at 37 °C with 5% CO_2_.

### Protein expression and purification

Codon-optimized DNA (Extended Data Table 2) encoding the ectodomain of VAR2CSA or TM284VAR1 was cloned into a pFastBac vector with the honeybee melittin signal peptide and a C-terminal 8×His tag. Baculoviruses were generated by the Bac-to-Bac system (Invitrogen), amplified using Sf21 insect cells and used to infect High Five cells for protein expression. Codon-optimized DNA encoding full-length DBLMSP or its DBL domain (residues 103-503) was cloned into a pcDNA vector with the IL-2 signal peptide and a C-terminal 8×His tag, and the resulting plasmids were transfected into HEK293F cells using polyethylenimine (PEI, Polysciences) for expression.

The cell cultures were collected by centrifugation at 500 × g, and the conditioned media were exchanged into binding buffer (25 mM Tris-HCl, pH 8.0, 150 mM NaCl). The recombinant proteins were isolated using the Ni-NTA affinity method and eluted using elution buffer (25 mM Tris-HCl, pH 8.0, 150 mM NaCl, 500 mM imidazole). They were further purified using gel filtration chromatography (Superdex 6 increase for VAR2CSA, TM284VAR1, and full-length DBLMSP; Superdex 200 increase for DBLMSP_DBL_) and eluted using binding buffer. The expression and purification of Fcμ-J were carried out as previously described ^4^. To obtain the complexes formed between the plasmodium proteins and Fcμ-J, purified plasmodium protein was mixed with Fcμ-J at a 2:1 molar ratio and incubated on ice for 1 h. The resulting complexes were then isolated using a Superdex 6 increase column in the final buffer (25 mM HEPES, pH 7.4, 150 mM NaCl).

To produce anti-CD20 or anti-RBD IgM molecules, heavy chain DNAs of the antigen-binding fragments of rituximab or BD-368-2 ^44^ were installed upstream of Fcμ (Data S1) in the pcDNA vector. The resulting anti-CD20 or anti-RBD heavy chain plasmids were transfected into HEK293F cells together with the corresponding light chain and J-chain expression plasmids using a 1:1:3 ratio. A C-terminal 8×His tag was added to the J-chain. The anti-CD20 or anti-RBD IgM proteins were then isolated from the conditioned medium using the Ni-NTA affinity method followed by a gel filtration step using the Superdex 6 increase column as described above. The buffer used for the gel filtration step was 25 mM Tris-HCl, pH 8.0, and 150 mM NaCl.

### Surface plasmon resonance (SPR)

SPR experiments were performed using a Biacore T200 (GE Healthcare). For this purpose, 200-300 resonance units (RU) of VAR2CSA or TM284var1, or 1800-2000 RU of DBLMSP were captured on a Series S Sensor CM5 Chip (Cytiva) in 0.01 M HEPES, pH 7.4, 0.005% (v/v) P20. Serial dilutions of purified Fcμ-J were then injected, ranging in concentration from 40 nM to 2.5 nM (2-fold dilutions). The SPR results were analyzed with Biacore Evaluation Software and fitted using a 1:1 binding model.

### Cryo-EM sample preparation, data collection and data processing

After gel filtration purification, purified ternary complexes containing Fcμ-J and the plasmodium proteins were concentrated to 0.9 mg/ml. These samples were then cross-linked with 0.05% glutaraldehyde (Sigma) for 10 min at 20 °C before the reactions were terminated by the addition of 1 M Tris-HCl (pH 7.4) to a final concentration of 100 mM. The cryo-grids were prepared using a Vitrobot Mark IV (FEI). The cross-linked samples were applied onto glow-discharged holey carbon gold grids (Quantifoil, R1.2/1.3) using a Vitrobot (FEI) at 4 °C with 100% humidity. The blotting time was 0.5~1.5 s, followed by a waiting time of 5 s. The grids were then plunged into liquid ethane. Grid screening was performed using a 200 kV Talos Arctica microscope equipped with a Ceta camera (Thermo Fisher/FEI). Good grids were recovered, and data collection was performed using a 300 kV Titan Krios electron microscope (Thermo Fisher/FEI) with a K3 direct detection camera. Statistics for data collection are summarized in Extended Data Table 1.

The cryo-EM data were processed following the workflows presented in Extended Data Fig. 2 and S3. Raw movie frames were aligned and averaged into motion-corrected summed images using MotionCor2 ^45^. The contrast transfer function (CTF) parameters were estimated using Gctf ^46^. The subsequent data processing was carried out with cryoSPARC ^47^ or RELION ^48^. Low-quality micrographs were removed manually, and particles were autopicked by template picking. All particles were subjected to several rounds of 2D and 3D classifications to exclude inaccurate particles. The favorite classes were selected for 3D refinement or mask-based local refinement focusing on protein interaction interfaces. The local resolution map was analyzed using ResMap ^49^ and displayed using UCSF ChimeraX ^50^.

### Structure building and refinement

The cryo-EM structure of Fcμ-J (PDB ID:6KXS) ^4^, the crystal structure of VAR2CSA_DBL3X_ (PDB ID: 3CML) ^51^, and the cryo-EM structures of the ID2a-ID2b and DBL4ε-DBL5ε regions of VAR2CSA (PDB ID: 7JGE, 7JGF) ^22^ were docked into the EM map of the VAR2CSA-Fcμ-J complex using UCSF Chimera ^50^ and then adjusted using Coot ^52^. Structural models of TM284VAR1_DBL4ζ_ and DBLMSP_DBL_ were first predicted using the tFold server (https://drug.ai.tencent.com/console/en/tfold) and then fitted into EM maps and adjusted using Coot. Structural refinement was performed using real-space refinement in Phenix ^53^.

### Strep pull-down assay

A twin-strep tag is present on Fcμ. For the pull-down experiments, 80 μg of purified plasmodium proteins and 40 μg of Fcμ-J proteins were incubated with StrepTactin beads (Smart Lifesciences) in binding buffer on ice for 1 h. The beads were spun down and then washed three times using binding buffer. Proteins retained on the beads were eluted using binding buffer supplemented with 10 mM desthiobiotin. The results were analyzed by SDS-PAGE and Coomassie staining.

### CDC assay

The complement dependent cytotoxicity (CDC) assay was performed using OCI-Ly10 cells that express CD20. Anti-CD20 IgM (6 nM) was incubated with serially diluted plasmodium proteins in 50 μl RPMI-1640 for 20 min and then mixed with equal volumes of OCI-Ly10 culture (~20,000 cells) and normal human serum complement (1:12.5 dilution, Quidel). The resulting mixtures were then transferred into a 96-microwell plate. After 6 h of incubation at 37 °C, 50 μl of CellTiter-Glo reagent (Promega) was added to each well and incubated for 10 min at room temperature. Luminescence was measured using a Cytation 5 cell imaging multimode reader (BioTek). The cell viability rate was analyzed according to the relative luminescence units and plotted against the plasmodium protein concentration using a 4-parameter curve-fit in GraphPad Prism.

### Data and Code Availability

Cryo-EM density maps of VAR2CSA-Fcμ-J, TM284VAR1-Fcμ-J, DBLMSP_DBL_-Fcμ-J, and DBLMSP2_DBL_-Fcμ-J have been deposited in the Electron Microscopy Data Bank with accession codes EMD-33542, EMD-33547 (EMD-33548 for the local map), EMD-33538 (EMD-33539 for the local map), and EMD-33805 (EMD-33806 for the local map), respectively. Structural coordinates have been deposited in the Protein Data Bank with accession codes 7Y0H, 7Y0J, 7Y09, and 7YG2.

## Acknowledgments

We thank the Core Facilities at the School of Life Sciences, Peking University for help with negative-staining EM; the Cryo-EM Platform of Peking University for help with data collection; the High-performance Computing Platform of Peking University for help with computation. We also thank the National Center for Protein Sciences at Peking University for assistance with the Biacore and BioTek Cytation Reader. Special thanks to J. Guo for insightful discussions on the CDC experiment. The work was supported partly by the Qidong-SLS Innovation Fund.

## Authors Contributions

J.X. conceived and supervised the project. C.J. and H.S. carried out most of the experiments with the help of C.S., Y.L., and S.C. T.H.S. provided the structural model of the IgM-C1 complex. J.X. wrote the manuscript, with inputs from all authors.

---

**Extended Data Fig. 1.**
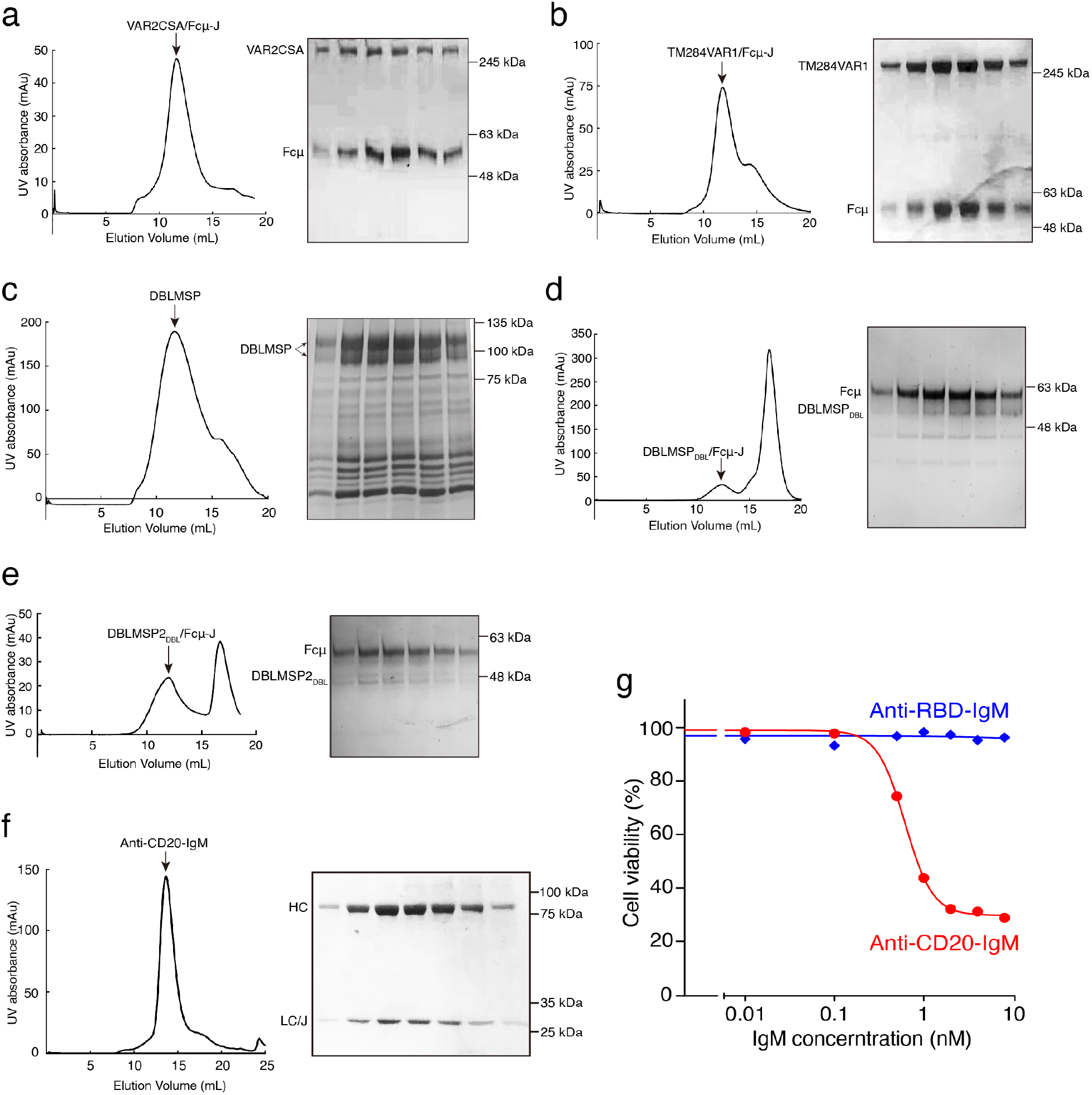
Purification and validation of protein samples used in this study. **a.** Size exclusion chromatography of the VAR2CSA-Fcμ-J complex (left) and SDS-PAGE analyses (right). **b.** Size exclusion chromatography of the TM284VAR1-Fcμ-J complex and SDS-PAGE. **c.** Size exclusion chromatography of the DBLMSP and SDS-PAGE. **d.** Size exclusion chromatography of the DBLMSP_DBL_-Fcμ-J complex and SDS-PAGE. **e.** Size exclusion chromatography of the DBLMSP_DBL_-Fcμ-J complex and SDS-PAGE. **f.** Size exclusion chromatography of purified anti-CD20 IgM and SDS-PAGE. HC: heavy chain; LC: light chain; J: J–chain. The light chain and the J-chain have similar molecular weights and could not be separated on the gel. **g.** Anti-CD20 IgM, but not another similarly recombinant IgM targeting the receptor binding domain of the SARS-CoV-2 spike (anti-RBD IgM), triggers CDC of OCI-Ly10 cells in the presence of human complement.

**Extended Data Fig. 2.**
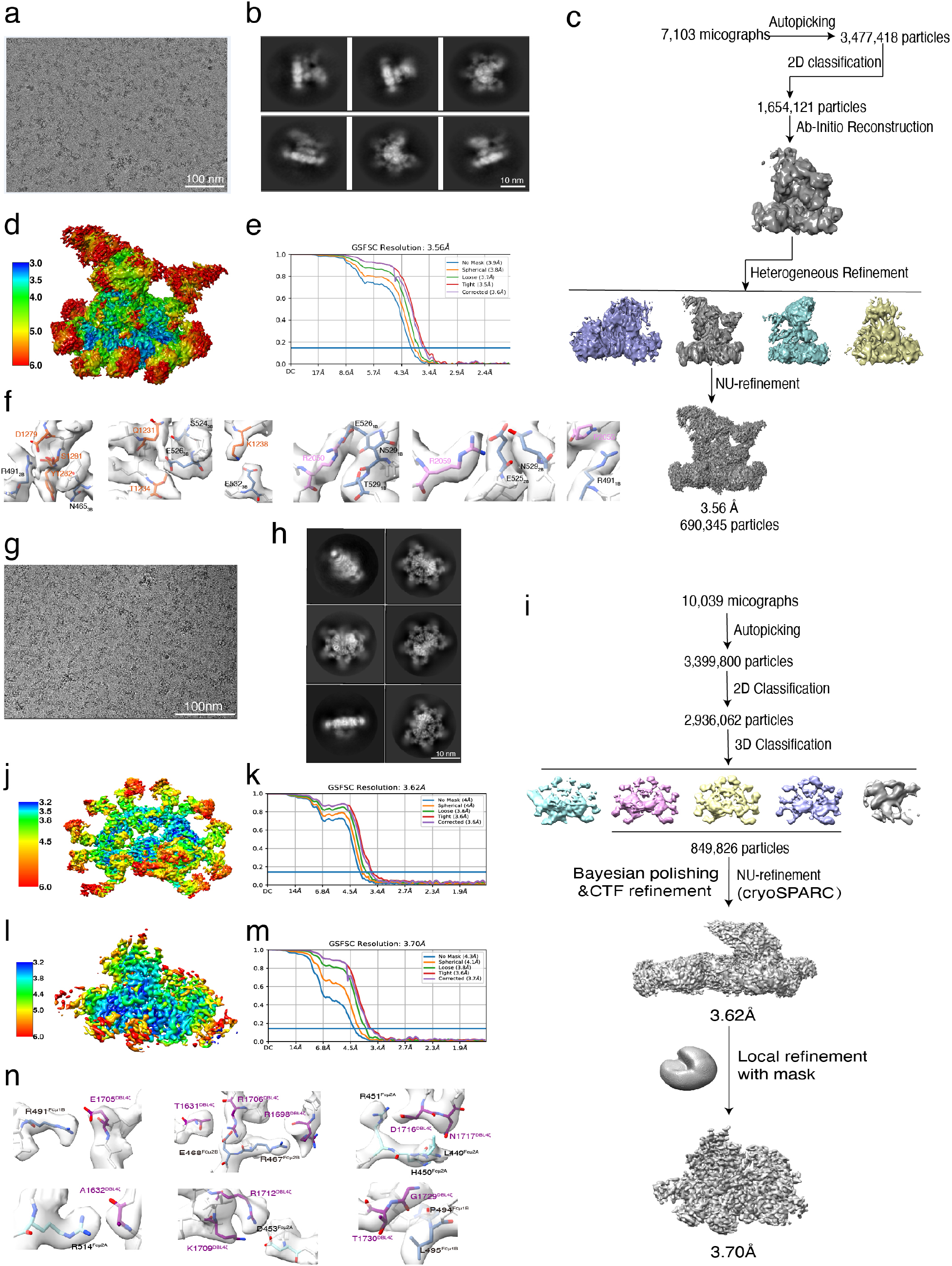
Workflow for the 3D reconstruction of the VAR2CSA-Fcμ-J and TM284VAR1-Fcμ-J cryo-EM structures. **a.** A representative raw cryo-EM image of the VAR2CSA-Fcμ-J complex. **b.** Representative 2D classes of VAR2CSA-Fcμ-J. **c.** Flow chart for image processing of VAR2CSA-Fcμ-J. **d.** Resolution estimations of the final map of VAR2CSA-Fcμ-J. **e.** Gold standard Fourier shell correlation (FSC) curves with estimated resolutions of VAR2CSA-Fcμ-J. **f.** Density maps of representative regions at the VAR2CSA-Fcμ interfaces. **g.** A representative raw cryo-EM image of TM284VAR1-Fcμ-J. **h.** Representative 2D classes of TM284VAR1-Fcμ-J. **i.** Flow chart for image processing of TM284VAR1-Fcμ-J. **j.** Resolution estimations of the final map of TM284VAR1-Fcμ-J. **k.** FSC curves with estimated resolutions of TM284VAR1-Fcμ-J. **l.** Resolution estimations of the local map around the binding interface of TM284VAR1-Fcμ-J. **m.** FSC curves for the local map of TM284VAR1-Fcμ-J. **n.** Density maps of representative regions at the TM284VAR1-Fcμ interface.

**Extended Data Fig. 3.**
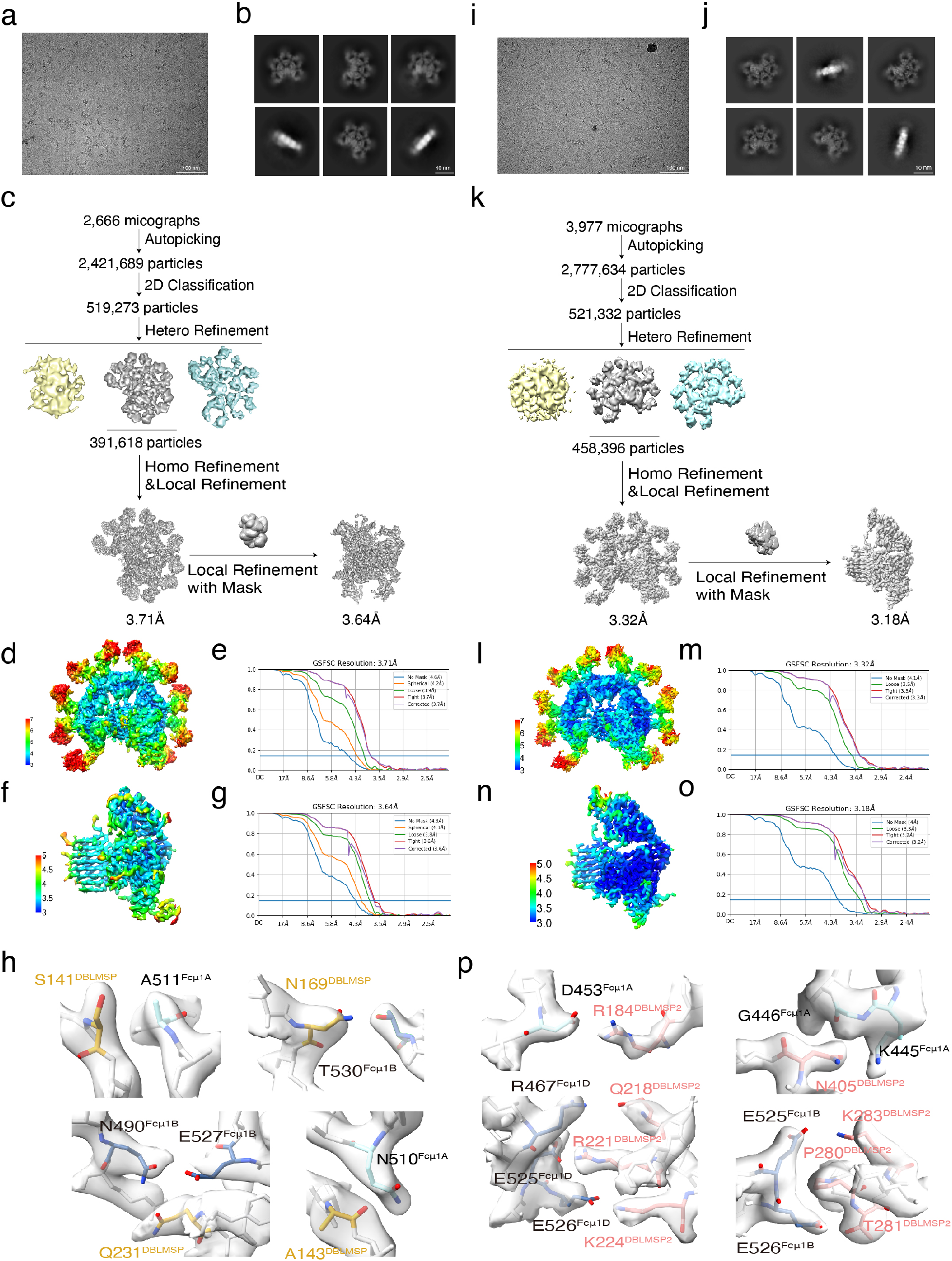
Workflow for the 3D reconstruction of the DBLMSP_DBL_-Fcμ-J and DBLMSP2_DBL_-Fcμ-J cryo-EM structures. **a.** A representative raw cryo-EM image of the DBLMSPDBL-Fcμ-J complex. **b.** Representative 2D classes of DBLMSPDBL-Fcμ-J. **c.** Flow chart for image processing of DBLMSPDBL-Fcμ-J. **d.** Resolution estimations of the final map of DBLMSPDBL-Fcμ-J. **e.** FSC curves with estimated resolutions of DBLMSPDBL-Fcμ-J. **f.** Resolution estimations of the local map around the binding interface of DBLMSPDBL-Fcμ-J. **g.** FSC curves for the local map of DBLMSPDBL-Fcμ-J. **h.** Density maps of representative regions at the DBLMSPDBL-Fcμ interface of DBLMSPDBL-Fcμ-J. **i.** A representative raw cryo-EM image of DBLMSP2DBL-Fcμ-J. **j.** Representative 2D classes of DBLMSP2DBL-Fcμ-J. **k.** Flow chart for image processing of DBLMSP2DBL-Fcμ-J. **l.** Resolution estimations of the final map of DBLMSP2DBL-Fcμ-J. **m.** FSC curves with estimated resolutions of DBLMSP2DBL-Fcμ-J. **n.** Resolution estimations of the local map around the binding interface of DBLMSP2DBL-Fcμ-J. **o.** FSC curves for the local map of DBLMSP2DBL-Fcμ-J. **p.** Density maps of representative regions at the DBLMSPDBL-Fcμ interface of DBLMSP2DBL-Fcμ-J.

**Extended Data Fig. 4.**
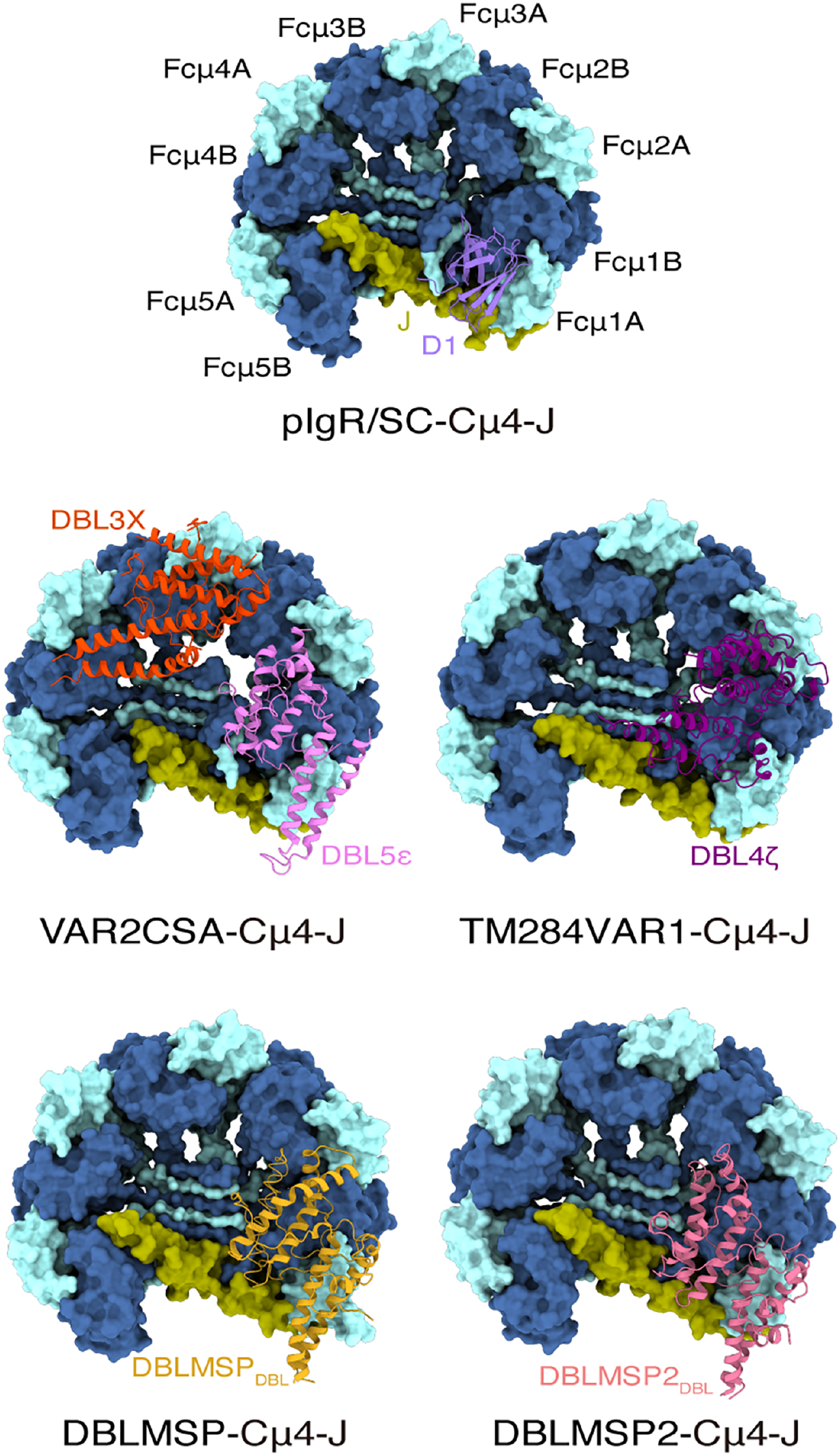
The binding sites of VAR2CSA, TM284VAR1, DBLMSP, and DBLMSP2 on Fcμ-J overlap with those of pIgR/SC. The structures of Fcμ-J in complexes with the D1 domain of pIgR/SC, VAR2CSA_DBL3X-DBL5ε_, TM284VAR1_DBL4ζ_, DBLMSP_DBL_, and DBLMSP_DBL_ are shown. Only the Cμ4 domains of Fcμ are depicted.

**Extended Data Fig. 5.**
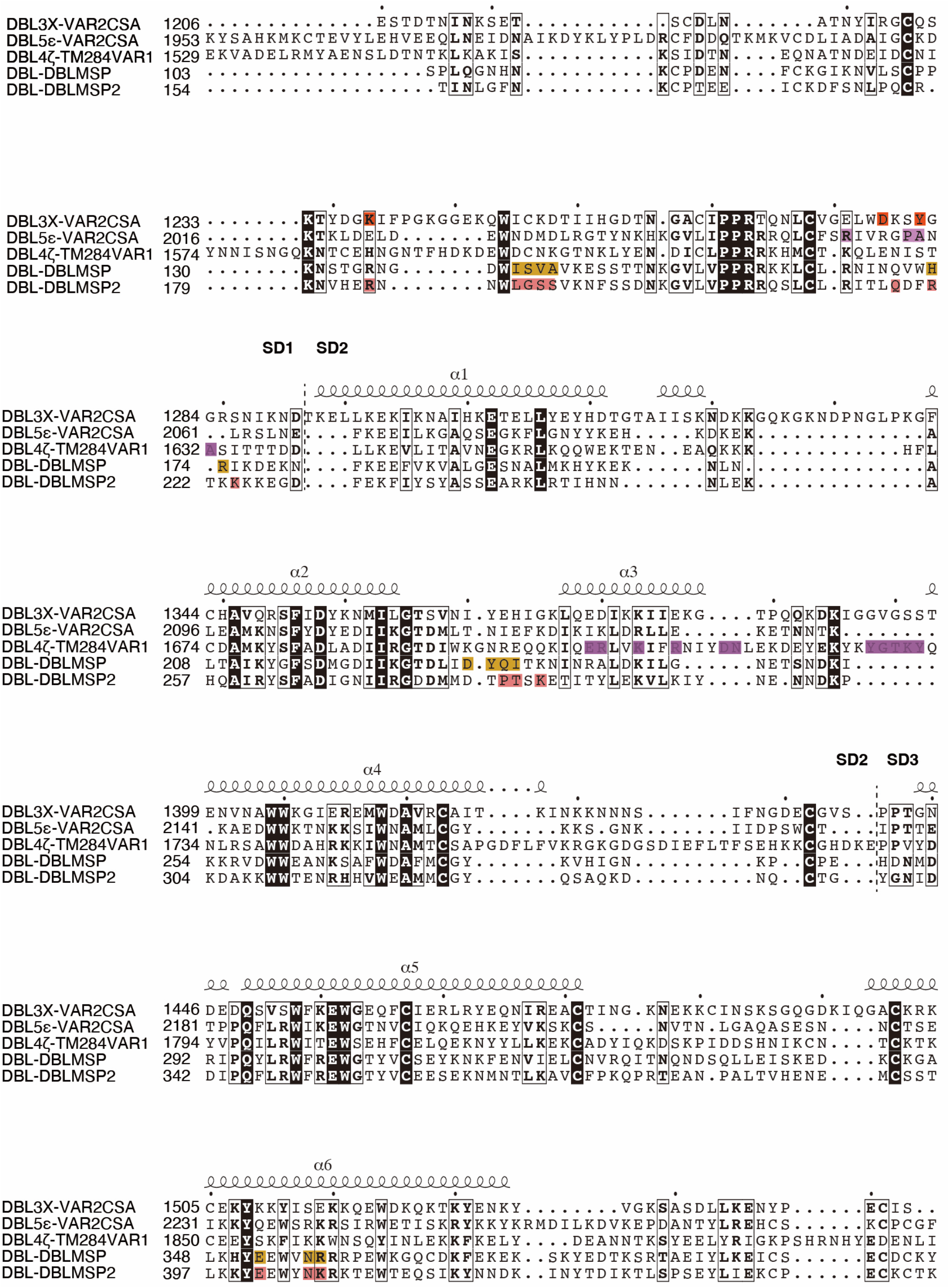
Sequence alignment of VAR2CSA-DBL3X, VAR2CSA-DBL5ε, TM284VAR1-DBL4ζ, DBLMSP-DBL, and DBLMSP2-DBL. Residues critically involved in bindingto IgM in each DBL domain are highlighted.

**Extended Data Fig. 6.**
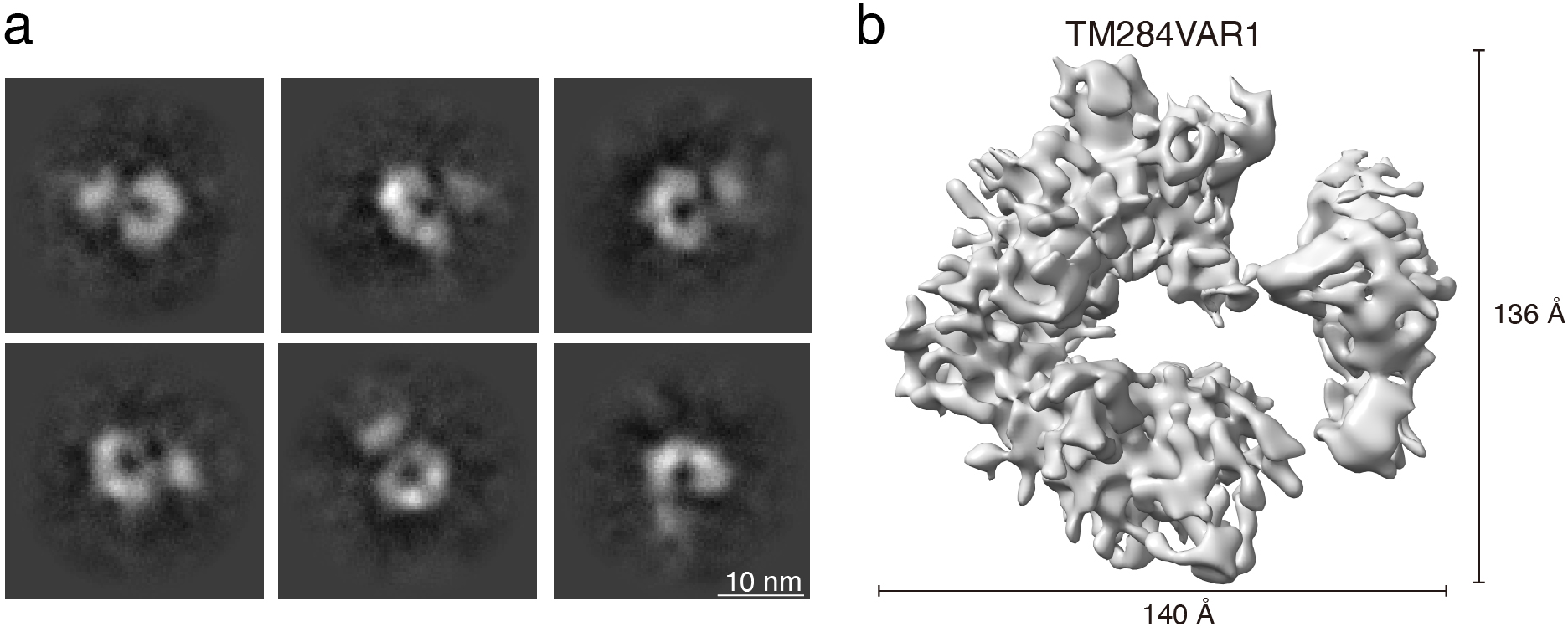
Cryo-EM reveals an elongated but flexible structural architecture of TM284VAR1. **a.** The 2D classification of the TM284VAR1 cryo-EM data suggests that it has a flexible structure. **b.** The 3D reconstruction of TM284VAR1 at ~8 Å.

**Extended Data Fig. 7.**
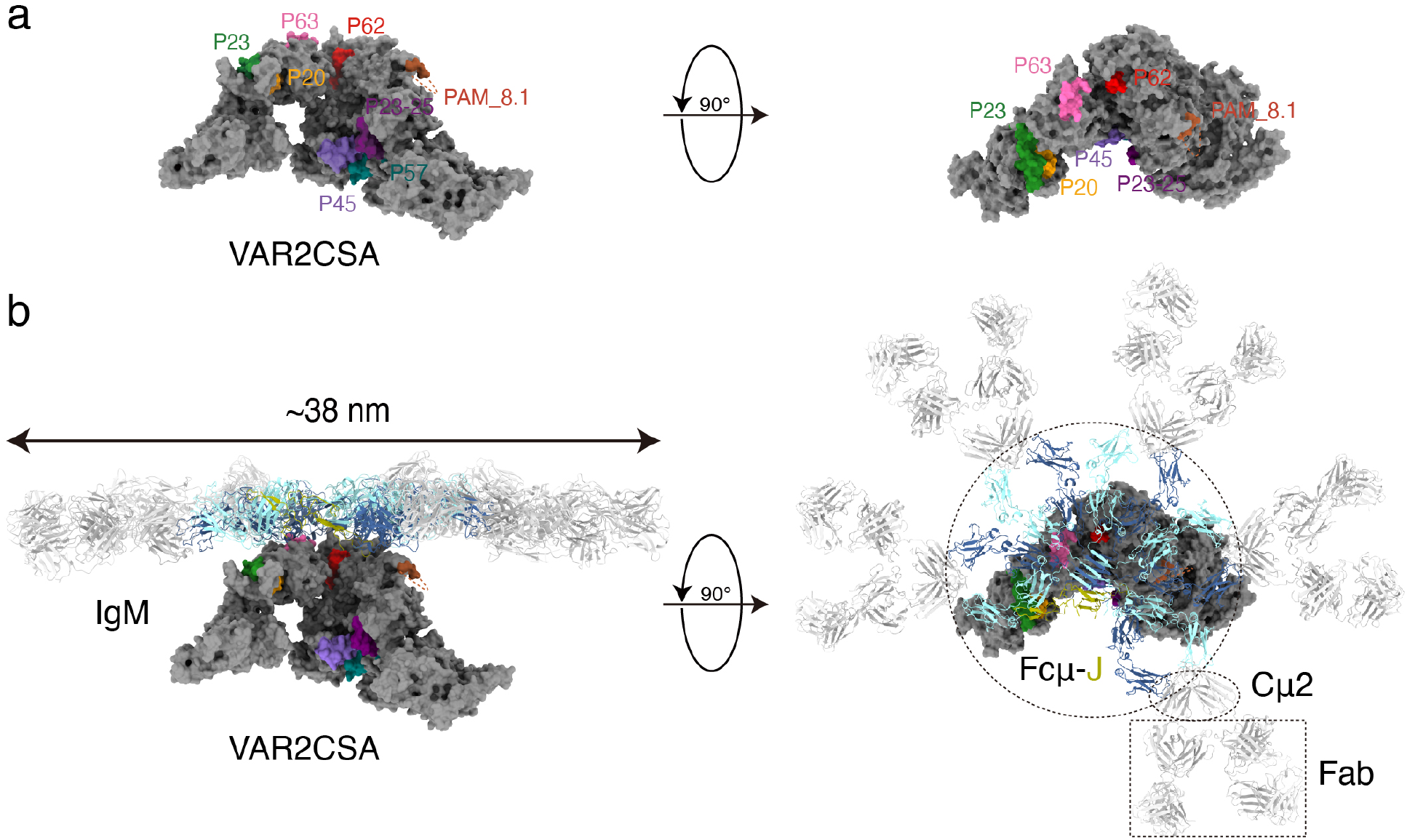
IgM masks IgG epitopes of VAR2CSA. **a.** Surface view of VAR2CSA, with the known IgG epitopes highlighted. **b.** A composite model of VAR2CSA in complex with a full-length IgM molecule.

**Extended Data Table 1.** Cryo-EM data collection, refinement and validation statistics.

|  | VAR2CSA-Fcμ-J | TM284VAR-Fcμ-J | DBLMSP-Fcμ-J | DBLMSP2-Fcμ-J |
| --- | --- | --- | --- | --- |
| **Data collection and processing** |  |  |  |  |
| Voltage (kV) | 300 | 300 | 300 | 300 |
| Microscope | FEI Titan Krios G3 | FEI Titan Krios G3 | FEI Titan Krios G3i | FEI Titan Krios G3 |
| Camera | K3 Summit (Gatan) | K2 Summit (Gatan) | K3 Summit (Gatan) | K3 Summit (Gatan) |
| Magnification (calibrated) | 81,000× | 165,000× | 64,000× | 81,000× |
| Electron exposure (e<sup>-</sup>/Å<sup>2</sup>) | 60 | 59.74 | 50 | 60 |
| Exposure rate (e<sup>-</sup>/Å<sup>2</sup>/s) | 18.75 | 11.688 | 19.76 | 21.47 |
| Number of frames collected per micrograph | 32 | 32 | 32 | 40 |
| Energy filter slit width | 20 eV | 20 eV | 20 eV | 20 eV |
| Automation software | EPU | SerialEM | EPU | EPU |
| Defocus range (μm) | -1.0 to -2.0 | -1.0 to -2.0 | -1.0 to -1.5 | -1.1 to -1.5 |
| Pixel size (Å) | 1.07 | 0.828 | 1.08 | 1.07 |
| Micrographs used | 7,103 | 10,039 | 2,666 | 3977 |
| Symmetry imposed | C1 | C1 | C1 | C1 |
| Initial particle images | 3,477,418 | 3,399,800 | 2,421,689 | 2777634 |
| Final particle images | 690,345 | 849,826 | 391,618 | 458396 |
| Overall map resolution at 0.143 FSC of masked reconstruction (Å) | 3.56 | 3.62 | 3.71 | 3.32 |
| Local map resolution at 0.143 FSC of masked reconstruction (Å) | / | 3.7 | 3.64 | 3.18 |
| Overall map sharpening B factor (Å<sup>2</sup>) | 138.3 | 163.7 | 105.7 | 90.6 |
| Local map sharpening B factor (Å<sup>2</sup>) | / | 124.7 | 116.5 | 89.5 |
| **Refinement** |  |  |  |  |
| Refinement package | Phenix v1.19 | Phenix v1.19 | Phenix v1.19 | Phenix v1.19 |
| Map-model CC |  |  |  |  |
| CC_mask | 0.77 | 0.73 | 0.78 | 0.75 |
| CC_box | 0.72 | 0.69 | 0.77 | 0.73 |
| CC_peaks | 0.63 | 0.58 | 0.68 | 0.66 |
| CC_volume | 0.75 | 0.7 | 0.76 | 0.74 |
| Model composition |  |  |  |  |
| Non-hydrogen atoms | 26,471 | 20,717 | 20,848 | 20859 |
| Protein residues | 3,336 | 2,616 | 2,639 | 2639 |
| Ligands | 12 | 12 | 12 | 12 |
| Bond lengths (Å) | 0.003 | 0.003 | 0.003 | 0.003 |
| Bond angles (°) | 0.627 | 0.687 | 0.674 | 0.697 |
| B factors (Å<sup>2</sup>) |  |  |  |  |
| Protein | 112.83 | 129.55 | 117.39 | 103.17 |
| Ligands | 77.77 | 93.52 | 87.04 | 69.23 |
| Validation |  |  |  |  |
| MolProbity score | 3.17 | 1.94 | 2.02 | 2.01 |
| Clashscore | 13.58 | 12.33 | 16.41 | 15.26 |
| Poor rotamers (%) | 0 | 0 | 0.3 | 0.04 |
| Ramachandran plot |  |  |  |  |
| Favored (%) | 95.54 | 95.1 | 95.66 | 95.31 |
| Allowed (%) | 4.46 | 4.9 | 4.34 | 4.58 |
| Disallowed (%) | 0 | 0 | 0 | 0.12 |
| Cβ outliers (%) | 0 | 0 | 0 | 0 |
| CaBLAM outliers (%) | 2.25 | 2.54 | 2.26 | 2.58 |

